# Time-delay model of perceptual decision making in cortical networks

**DOI:** 10.1101/348276

**Authors:** Natalia Z. Bielczyk, Katarzyna Piskała, Martyna Płomecka, Piotr Radziński, Lara Todorova, Urszula Foryś

**Affiliations:** Stichting Solaris Onderzoek en Ontwikkeling, Nijmegen, the Netherlands; Faculty of Mathematics, University of Warsaw, Warsaw, Poland; Faculty of Social Sciences, Radboud University Nijmegen, Nijmegen, The Netherlands; Laboratory of Brain Imaging, Neurobiology Center, Nencki Institute of Experimental Biology of Polish Academy of Sciences, Warsaw, Poland; Donders Centre for Cognitive Neuroimaging, Donders Institute for Brain, Cognition and Behavior, Nijmegen, the Netherlands

**Keywords:** perceptual decision making, Hopf bifurcation, perceptual switches

## Abstract

It is known that cortical networks operate on the edge of instability, in which oscillations can appear. However, the influence of this dynamic regime on performance in decision making, is not well understood. In this work, we propose a population model of decision making based on a winner-take-all mechanism. Using this model, we demonstrate that local slow inhibition within the competing neuronal populations can lead to Hopf bifurcation. At the edge of instability, the system exhibits ambiguity in the decision making, which can account for the perceptual switches observed in human experiments. We further validate this model with fMRI datasets from an experiment on semantic priming in perception of ambivalent (male versus female) faces. We demonstrate that the model can correctly predict the drop in the variance of the BOLD within the Superior Parietal Area and Inferior Parietal Area while watching ambiguous visual stimuli.

**Author summary:** Human cortex is a complex structure composed of thousands of tangled neural circuits. These circuits exhibit multiple modes of activity, depending on the local balance between excitatory and inhibitory activity. In particular, these circuits can exhibit oscillatory behavior, which is believed to be a manifestation of a so-called criticality: balancing on the edge between stable and unstable dynamics. Circuits in the cortex are responsible for higher cognitive functions such as, in example, perceptual decision making, i.e., evaluating properties of objects appearing in the visual field. However, it is not well known how aforementioned balancing on the edge of instability influences perceptual decision making.

In this work, we build a model to simulate dynamics of a very simple decision-making network consisting of two subpopulations. We then demonstrate that criticality in the network can account for ambiguity in decision making, and cause perceptual switches observed in human experiments. We further validate our model with datasets coming from a functional Magnetic Resonance Imaging experiment on semantic priming in perception of ambivalent (male versus female) faces. We demonstrate that the model can correctly predict the drop in the variance of the BOLD within the parietal areas of the cortex while watching ambiguous visual stimuli.

## Introduction

### Models of perceptual decision making

Decision making – from simple perceptual choices such as deciding upon the spatial orientation of the Necker cube [1] to complex choices such as choosing the best vacation destination – is one of the basic functionalities of the human brain [2]. While making a decision, one needs to evaluate the options, and withdraw all but one. The concept of such a winner-takes-all mechanism is well rooted in computational neuroscience. In general, the process of evaluating 7 the options is energy-costly as it requires assessing the utility of each option [3]. For instance, choosing between an apple and an orange requires evaluating the taste, smell, nutritious value, appearance, price and other potentially useful features. In models of decision making, this assessment is often envisaged from the Bayesian point of view: the brain as a machine collecting evidence on behalf of one or the other option. According to this viewpoint, the brain needs time to aggregate enough evidence favoring one alternative over the other before reaching the critical amount of evidence that allows for making the decision. This is a time-costly process, however taking time helps in isolating the relevant sensory information from the noise and therefore to optimize the response [4].

In this work, we focus on modeling the most basic, perceptual decision making in neuronal networks. In such conditions, the network needs to disambiguate between two sensory stimuli. In laboratory conditions, this phenomenon is usually investigated with use of two-alternative forced choice experiments [5]. In the famous study, Michael Shadlen trained rhesus monkeys [6] to distinguish between the dots moving on the screen towards either right or left direction. In each trial, the stimulus presented on the screen was ambiguous: multiple dots were moving at the same time, and the task for the monkey was to estimate in which direction the *majority* of dots were moving. It was found that the monkeys followed the psychometric curve: they were most uncertain about the direction once the stimulus was the most ambiguous (almost 50% of dots in each direction). Moreover, the response of the Lateral intraparietal cortex (LIP area) was substantially delayed with respect to the visual stimulus, which supports the hypothesis on the aggregation of sensory information over time in the decision making networks [7, 8].

In the literature, one model proposed to study the perceptual decision making in neuronal networks is the slow reverberation mechanism by Wang [9], which was designed to model the Shadlen’s experiments on rhesus monkeys. In this model, two populations of densely interconnected, spiking neurons compete with each other when supplied by noisy inputs. In this model, the binary decision is delayed by 1 – 2[s] with respect to the onset of the stimulus because of the recurrent excitation mediated by the slow NMDA receptors. Furthermore, the network is prone to making incorrect decisions in a low signal-to-noise ratio regime (when a stimulus has a low magnitude as compared to the magnitude of the background noise). The success rate in a function of the stimulus reproduces the psychometric curve previously found in Shadlen’s experiments.

### The brain at the edge of instability

Second important paradigm with respect to the dynamics of cortical networks, relates to neuronal oscillations as a result of the fact that, in terms of dynamical systems, the brain occupies a state at the edge of instability.

*Criticality* [10] is a concept that refers to all neuronal networks that, as dynamical systems, might operate nearby critical points. Biological systems have a tendency to spontaneously reorganize to a critical point between order and chaos [11] which is referred to as a self-organized criticality. For instance, intrinsic oscillations found in EEG studies are believed to be an evidence for self-organized criticality in brain networks (more precisely, as evidence for brain networks keeping balance around the Hopf bifurcation [12]). But how do the oscillations arise? Some hint was given by Deco et al. [13] who demonstrated in a simulation study that a network of nodes with sigmoidal transfer functions, coupled with transmission delays, can start oscillating in the gamma range when tuned to a subcritical state.

However, although criticality in cortical networks has been researched, *decision making around the critical point* is still an unexplored topic. In this work, we investigate how dynamic states close to the critical points can account for ambiguity in decision making.

### Model of ambivalence in perceptual decision making

Since Wang’s work was published, multiple rodent [14] and computational [15] models were proposed to study perceptual choices. Most of these models aim to reproduce the psychometric curve within the signal detection theory framework [14]. In particular, few models were developed to study perceptual rivalry as a winner-takes-all mechanism in neural mass models with mutual inhibition and additionally: with balance between noise and adaptation strength in the networks [16], with a solution to the Levelt’s fourth proposition [17], with slow negative feedback in the mutual inhibition in the form of spike frequency adaptation [18], or with short short-term memory in the mutual inhibition [19].

In this work, we investigate a winner-take-all population model with delayed local inhibition to characterize perceptual switches in the binary decision making. We have proposed this model in our previous work [20]. In this work, we investigated mathematical properties of this model, especially with respect to the influence of the value of inhibitory delays on the emergence of temporal switches in the decision making.

In this paper, we first demonstrate that local inhibition within the competing neuronal populations can lead to a Hopf bifurcation. As a result, providing the network with a weak stimulus can induce ambivalent behavior, in which probabilities of making perceptual choices on behalf on both options vary over time which can lead to perceptual switches. In presence of a strong stimulus on the other hand, the probability of taking one of the possible decisions approaches 100% early on, which leads to an unambiguous decision on behalf of one option.

Then, we focus of further investigating the properties of this model, but this time, with respect to variables important for modeling biological systems. Namely, we are interested in dependence of the certainty in the binary decision making on the stimulus strength and the magnitude of background noise in the system. We demonstrate that this model reproduces psychommetric curves observed in human and animal experiments on perceptual decision making.

We also validate the model with experimental functional Magnetic Resonance Imaging (fMRI) datasets coming from an experiment on the impact of semantic priming on the perception of ambivalent (male versus female) faces. FMRI studies on the perceptual ambivalence in humans suggest that frontoparietal regions cope with visual ambiguities in a top-down fashion [21, 22]. Transcranial magnetic simulation (TMS) studies demonstrate that the Superior Parietal Lobe (SPL) controls switching attention between competing percepts [23-25]^1^. In general, SPL is involved in manipulating information within the working memory [28], including switching between bistable percepts. In addition, it has been found that Inferior Parietal Lobe (IPL) can contribute to the inference and decision making during perceptual ambiguity [22, 29]. However, IPL have not been proven to play a causal role in perceptual disambiguation.

Therefore, we assume that the decision making characterized by our model takes place in the SPL. The model predicts that in presence of ambivalent facial stimuli, the variance of the *joint* signal from the competing populations will drop with respect to the signal collected both in absence of the signal and in presence of a strong signal (representing unambiguous stimulus). We validate this prediction using datasets from an experiment on perception of ambiguous (male versus female) faces. We demonstrate that while viewing ambivalent faces, the variance of the BOLD response within the SPL and within IPL decreases as compared to the variance of the BOLD in a control condition and while watching unambiguous faces.

## Methods

### Model description

Let us consider two neuronal populations (1 and 2, Fig 1). Neurons in the first cluster project to the neurons in the second cluster with synaptic weight *w*_1_(*t*), while neurons in the second cluster project back to neurons in the first cluster with synaptic weight *w*_2_(*t*). As synapses are plastic; the synaptic weights evolve over time. Both neuronal populations receive self-inhibition with delays τ_1_, τ_2_. hat we are interested in here, is the slow, complex synaptic transmission processes, which can occur over periods of hundreds of milliseconds to minutes [30]. These slow transmission molecular pathways involve at least 100 compounds, biogenic amines, peptides, and amino acids. 109

**Figure 1.**
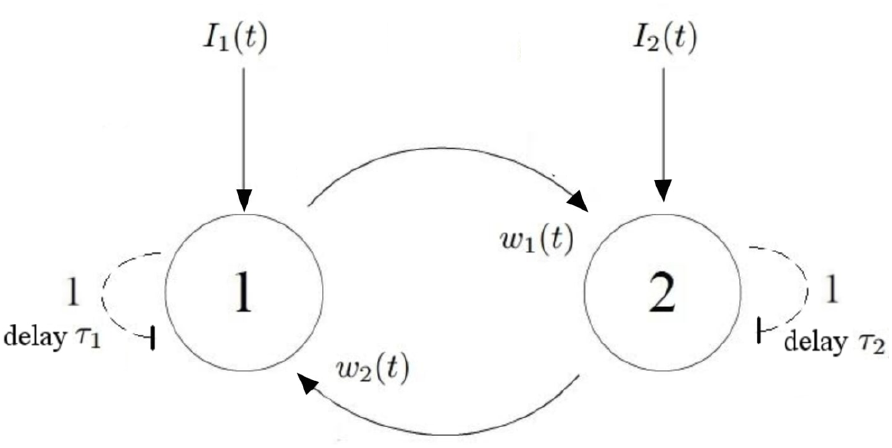
A population model of decision making. Neurons in the first cluster project to the neurons in the second cluster with synaptic weight *w*_1_(*t*), while neurons in the second cluster project to neurons in the first cluster with synaptic weight *w*_2_(*t*). Neurons receive external inputs *I*_1_(*t*) and *I*_2_(*t*), respectively. As this is a rate model, *I*_1_(*t*), *I*_2_(*t*) have a unit of [1/*s*]. In addition, both neuronal populations receive self inhibition with delays (dashed arrows with flat heads), *τ*_1_ and *τ*_2_ respectively. The dynamics of this system is described by Eq. 2.

Modeling of such systems with bilinear population models was introduced by Ermentrout et al. [31]):

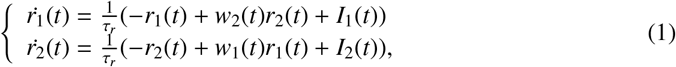

where *I*_1_(*t*), *I*_2_(*t*) denote inputs to the neuronal populations ([1/*s*]) and τ_*r*_is the time scale of firing rates ([*s*]).

In this work, we extend this model by adding self-inhibition delays to the populations [32] *τ*_1_, *τ*_2_ (Fig 1). This term is crucial for the dynamics, as it can lead to Hopf bifurcation^2^.

We also consider asymmetry coming from a perceptual stimulus *σ*(*t*). In our model, the stimulus *σ*(*t*) related to the perceptual stimulation targets the population 1 and is an addition to the input *I*_1_(*t*) in Eq. 2 (a constant pulse σ(*t*) = σ):

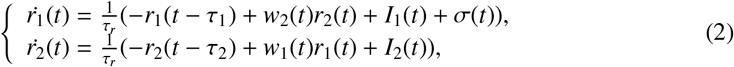

In general, the two competing neuronal populations are not an isolated system but are embedded in a bigger cortical network. Therefore, one can assume that these inputs represent the background activity within the system and are higher than zero even in resting state (in the absence of the experiment-related stimulus σ(*t*)). In our simulations, we assume that both the inputs *I*_1_(*t*), *I*_2_(*t*) are constant and equal, as they represent the resting state.

Furthermore, in our model, synapses are plastic:

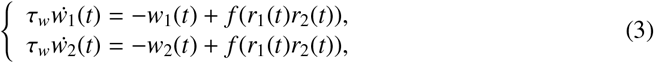

where 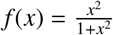 is a sigmoidal transfer function, a.k.a. Hill function [35], and *τ_w_* denote the time scales of synaptic weights ([*s*]). Further, we assume that synaptic weights *w_i_* adjust themselves to the changes in firing rates *r*_1_ and *r*_2_ instantaneously (*τ_w_* ≪ 1), which allows to use quasi-steady approximation for System (3) (*τ_w_w˙*_1_(*t*) ≈ *τ_w_w˙*_2_(*t*) 0).

As in the resting state (in the absence of external stimulus), this system receives equal inputs *I*(*t*) to both nodes, the system is perfectly symmetric and both populations will be firing with the same firing rate. Once one of the populations, i.e., population 1, receives external stimulation, the symmetry breaks down. Decision making in this system is associated with decoding *which* of the two populations received an additional stimulus (without estimation of the stimulus magnitude). The decoding is based on the ratio of the two firing rates: the higher the difference *r*_1_(*t*) – *r*_2_(*t*), the more likely the decision that the population 1 is the one to have received the stimulation. The evidence behind each of the two options accumulates over time (which reflects the Bayesian view at decision making mentioned in the Introduction). Therefore, the psychometric function takes the integral form of

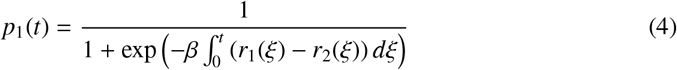

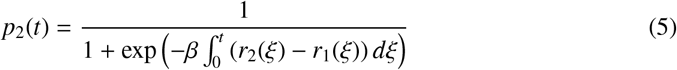

Values *p*_1_(*t*), *p*_2_(*t*) can only asymptotically approach 1, therefore we add a condition that the decision is made when at a given time point *t*, *p*(*t*) surpasses the threshold value of 0.99 for one of the populations. As population 1 receives the input, this is a correct decision if *p*_1_(*t*) > 0.99 and a wrong decision if *p*_2_(*t*) > 0.99. In this model, β is a parameter influencing the sigmoidal function of the decision probability with respect to the cumulative difference 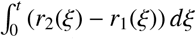.

In this work, we perform the stability analysis of the system 2- 3, and find a regime in which:

1. in absence of a stimulus, no decision can be made
2. in presence of a weak stimulus, we obtain perceptual ambivalence: *p*_1_(*t*) and *p*_2_(*t*) intersect with each other at different points in time
3. in presence of a strong stimulus, the network makes the right decision

We use Mathematica^®^(http://www.wolfram.com/mathematica/) to implement a system of delayed differential equations described by Eq. 2 and 3, in three regimes: the resting state, a weak stimulus regime and a strong stimulus regime. We then investigate the following aspects of the model:

#### Certainty of the decision making in a function of model parameters

A model for perceptual decision making should be able to reproduce the psychometric curve. Therefore, we define *certainty* of the decision making as

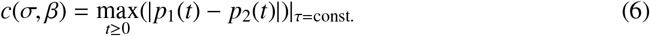

and investigate how this quantity depends on the stimulus magnitude *σ* and slope parameter *β*.

#### Influence of the stimulus duration on the decision making

We also explore how the duration of the stimulus influences the effect of perceptual ambivalence.

#### Influence of the background noise on the decision making

In the version of the model introduced above (Eq. 2), there is no stochasticity. However, as we model biological systems, there should be a smooth dependency between success rate in the decision making and the level of stochasticity in the model. Therefore, let us consider stochasticity in the neuronal dynamics. Here, we introduce the noise term as two independent Wiener processes controlled by parameter *b* as follows:

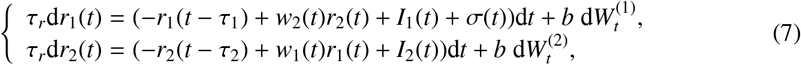

Then, we investigated the success rate in the decision making with respect to the parameter *b* (*σ* = 0.3, *τ* = 1.4). In order to be consistent with the fMRI experiment used for validation, we defined the success rate on the basis of decisions made at random time points in the range between 5 and 7 [s]. For each value of parameter *b*, we simulated 20, 000 iterations of the system on the interval 0-8 [s] in *R*.

Open source implementation of our model implemented in Mathematica (together with simulations of the noisy system in *R*) is available at https://github.com/PiotrRadzinski/DelayModelofPerceptualDecisionMaking.

### Validation in fMRI datasets

#### Prediction obtained from the model

Our model exhibits dynamics that demonstrates decisional ambivalence for low stimulus magnitudes *σ*. As the decision making neuronal populations are close in space, they cannot be distinguished from each other with use of macroscopic neuroimaging tools such as fMRI or EEG. However, as in the low signal magnitude, the model predicts that firing rates oscillate in antiphase, the *joint signal* coming from populations *r*_1_(*t*), *r*_2_(*t*) should have *low variance* when compared to the high signal magnitude regime.

On the basis of this reasoning, we made prediction upon the expected variance of the joint signal coming from the two populations in the system in presence of a very weak stimulus as lower than in strong stimulus regime. In fact, in functional Magnetic Resonance Imaging, the signal recorded in the experiment does not correspond the neuronal firing described by Eq. 2 directly, but rather, relates to the oxygen consumption by neurons during the firing. This is a weakly nonlinear dependency, where the BOLD response can be characterized as a linear convolution of the neuronal signal with a so-called hemodynamic response function [36] (explained in S2). However, as the BOLD signal is a linear convolution, the prediction of the model will hold.

We then confront this prediction with the experimental datasets coming from fMRI experiment on gender perception.

#### Subjects

Twenty-four female native Dutch speakers participated in the fMRI experiment and gained monetary compensation for their participation. Only female participants were recruited for the study, in order to avoid gender-related confounding factors [37]. The study was approved by the local ethics committee (CMO Arnhem-Nijmegen, Radboud University Medical Center, ethical approval for studies on healthy human subjects at the Donders Centre for Cognitive Neuroimaging, no *ECG*2012 – 0910 – 058) and conducted in accordance with their guidelines. All participants signed informed consent forms before the experiment. The data from seven subjects were excluded from the analysis: 3 subjects failed to finish the task and 4 subjects exhibited head motion that exceeded the maximum acceptance rate of 2[mm]. The remaining 17 subjects (females, age 18 − 29 years) reported no neurological diseases, and had normal or corrected-to-normal vision.

#### Stimuli

A set of realistic 3D faces was morphed across gender (from extremely female to extremely male) using FaceGen Modeller 3.5 (Singular Inversions, www.facegen.com, Fig 2 A). The morphing procedure started from 40 distinct faces. For each face, we gradually modulated gender features in 5 steps with the same amount of feature transformation in each step. The technical details upon the computation performed by the software are introduced in [38]. The face stimuli were presented frontally and cropped around the oval of the face. We controlled for luminance using SHINE toolbox for MATLAB [39]. The perceptual boundary within gender continuum of faces was established in a separate behavioral experiment.

**Figure 2.**
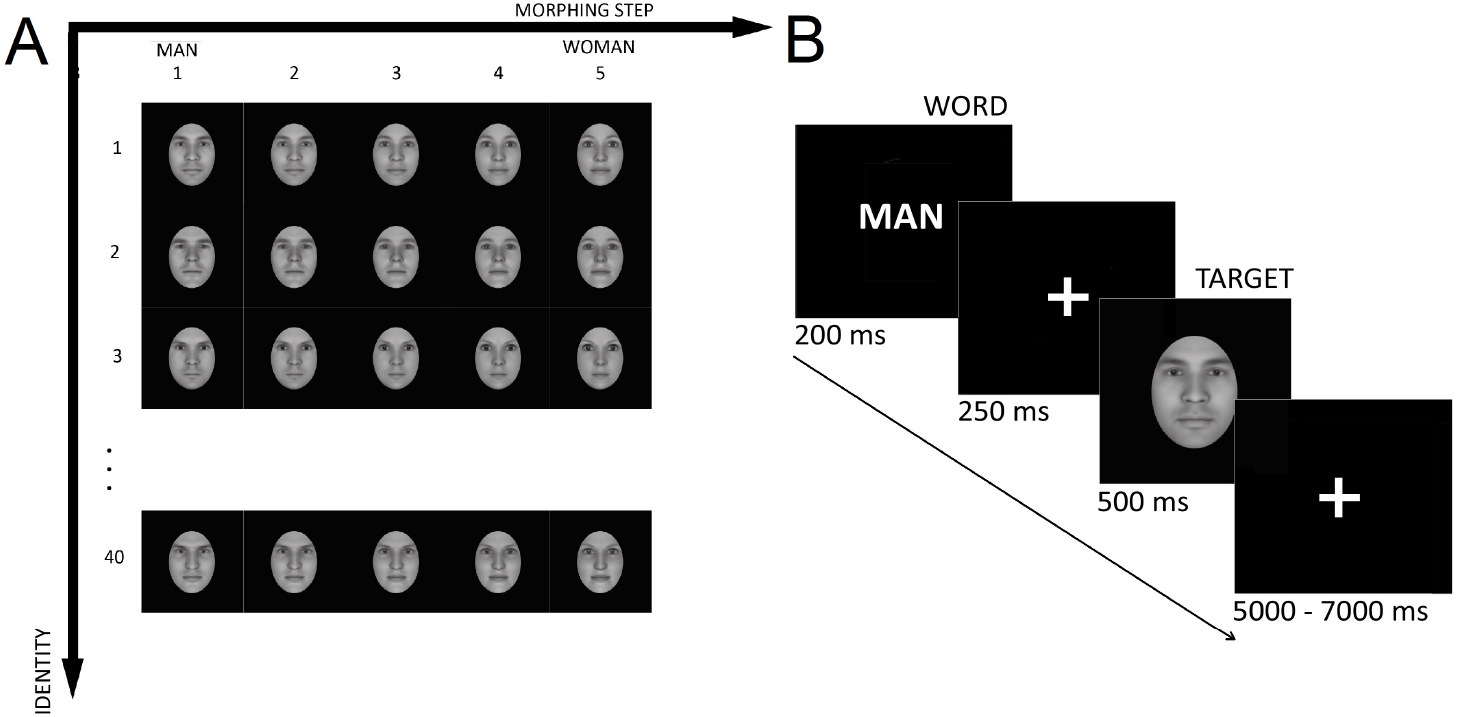
**A:** The set of stimuli. Each of the 40 distinct faces was morphed from male (1) to female (5). **B:** Experimental design. Each trial started with the word (*man or vrouw*) that was presented for 0.2[s]. After the fixation cross (0.25[s]), the face was briefly presented for the period of 0.5[s], followed by a randomized inter-stimulus interval (ISI) in the time range of 5 − 7[s].

#### Experimental Design

Each trial started with priming: presentation of a gender-related word (*man* or *vrouw*) for 0.2[s]. Then, after the fixation cross (0.25[s]), a face was presented (0.5[s]), followed by an inter-trial period of a randomized length of 5 – 7[s] (Fig 2 B). Participants were asked to perform a matching task: respond *yes* if a word and subsequent picture corresponded in gender, and *no* otherwise. The experiment was carried out in Dutch. The buttons were counterbalanced across subjects. The experiment was divided into 6 blocks in order to avoid fatigue. Each block consisted of 50 trials. The order of stimuli was randomized across blocks and participants. We used Presentation software (version 17.1, www.neurobs.com) in order to screen the stimuli during the experiment.

#### MR image acquisition

Functional images were acquired using 3T Skyra MRI system (Siemens Magnetom), T2* weighted echo-planar images (gradient-echo, repetition-time *TR* = 1760[ms], echo-time *TE* = 32[ms], 0.7[ms] echo spacing, 1626 hz/Px bandwidth, generalized auto-calibrating partially parallel acquisition (GRAPPA), acceleration factor 3, 32 channel brain receiver coil). In total, 78 axial slices were acquired (2.0[mm] thickness, 2.0 * 2.0[mm] in plane resolution, 212[mm] field of view (FOV) whole brain, anterior-to-posterior phase-encoding direction).

#### Preprocessing of functional MR imaging data

The data reprocessing was performed using SPM12 (Welcome Trust Center for Neuroimaging, University College London, UK). Functional scans were realigned to the first scan of the first run with further realignment to the mean scan. We performed slice-time correction on realigned images to account for differences in image acquisition between slices. Motion-related components were removed from the data using a data-driven ICA-AROMA [40]. Denoised functional scans were spatially normalized to the Montreal Neurological Institute (MNI) space [41, 42] without changing the voxel size. Normalized data were smoothed spatially with a Gaussian kernel of 6[mm] full-width at half-maximum.

#### Definition of Regions of Interest

We extracted region-of-interest (ROI) mask using Anatomical Automatic Labeling atlas (AAL, [43]). According to our a priori hypothesis, we preselected the bilateral SPL (4288 voxels) and the bilateral IPL (3792 voxels).

#### Computing variance of the BOLD

As the experiment only contained a few ambivalent facial stimuli, we performed a group analysis. Firstly, we extracted the BOLD within the bilateral SPL and IPL from each subject, and normalized the BOLD time series throughout the experiment to the mean of 0 and variance of 1 within each subject. Next, we extracted all the frames registered after the ambiguous stimuli were presented on the screen, and before the onset of the next stimulus. Regardless of the priming, we interpreted the pictures with morphing at stage 3 (the morphed images exactly halfway between male and female faces) as the ambiguous stimuli. We obtained 4,624 frames in total for this weak-stimulus condition. Lastly, we extracted all the frames registered after the faces of morphing phase 1, 2, 4 and 5 (unambiguous) were presented on the screen, and before the onset of the next stimulus. This resulted in 18,373 frames for the strong-stimulus condition. Given these two outcome vectors of BOLD values for each of the two ROIs, we performed post hoc pairwise F-tests to test for the difference in variance between the two regimes.

## Results

### Simulation results

#### Noiseless case

Full stability analysis of this model in the resting state (σ = 0), is given in S1. In the following section, we present exemplary simulation results. In our example, we assume that the inputs to the system during resting state are equal and constant (*I*_1_(*t*) = *I*_2_(*t*) = *I*(*t*) = 0.4[1/*s*]), and *β* = 1.

In absence of a stimulus, the dynamic system is launched from the reference point, and the two populations will exhibit identical dynamics, and therefore also identical firing rates in every point in time (regardless of the value of the synaptic delay *τ*). Therefore, the accumulated evidence *p*(*t*) in both nodes will be constant over time and equal to 0.5, as demonstrated in Fig 3 A.

**Figure 3.**
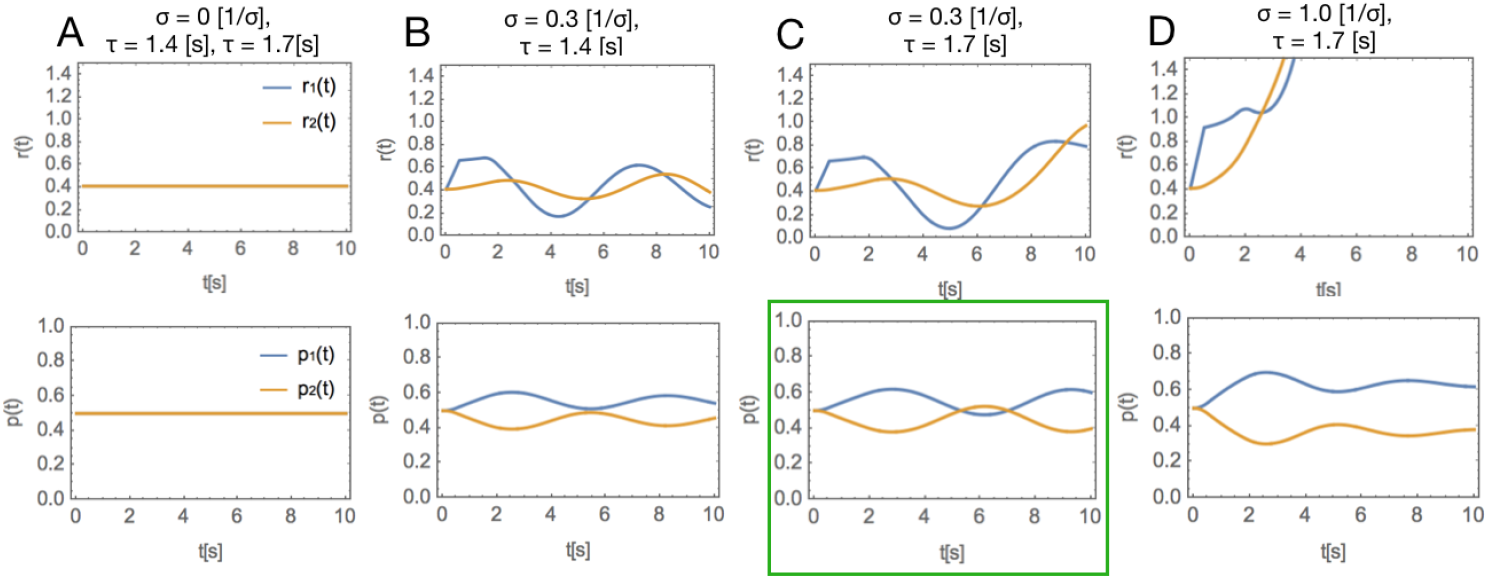
Dynamics in the system in a function of *τ* and *σ* (*β* ≡ 1, *I*_1_,(*t*) = *I*_2_,(*t*) ≡ 0:4). Upper row: the neuronal dynamics *r*(*t*). Lower row: accumulated evidence *p*(*t*). **A:** no stimulus. The two populations have a constant firing rate, and the associated cumulative evidence is stable in time and equal to 50%, for both *τ* = 1:4[*s*] and *τ* = 1:7[*s*]. Since there is no stimulus, from the symmetry of the system, we have *r*_1_(*t*) ≡ *r*_2_(*t*). **B:** a weak stimulus (*σ* = 0:3[1=*s*]), for the subcritical value of *τ* = 1:4[*s*]. The system dynamics is still stable, and perceptual ambivalence does not occur. C: a weak stimulus (*σ* = 0:3[1/s]), for the supercritical value of *τ*= 1:7[s]. In this regime, accumulated evidence behind choices 1 and 2 fluctuates over time, which may account for perceptual switches. **D:** a strong stimulus (*σ* = 1:0[1/*s*]) for supercritical value of *τ* = 1:7[*s*]. The effect of ambiguity disappears, and the cumulative evidence for choice 1 is higher than the cumulative evidence for choice 2 for the whole duration of the experiment.

In order to observe an interplay between the two populations, symmetry must be broken by adding a stimulus to one of the populations. In DDEs, it is convenient to perform this stimulation by adding a pulse σ to one of the populations. Thus, we set the value of the firing rate in population 2 to *r*_2_(*t*) = 0.4 and the value of the firing rate in population 1 to *r*_1_(*t*) = 0.4 + *σ*(*t*) on the time interval [0, *t_max_*]. In the example on Fig 3, *t_max_* = 0.5[*s*], which matches typical duration of presented visual cues in cognitive experiments on visual perception.

As demonstrated in Fig 3 B, in presence of a weak stimulus *σ* = 0.3[1/*s*] and delay *τ* below the critical point (*τ* = 1.4[*s*]), the system dynamics is still stable, and perceptual ambivalence does not occur. However, as demonstrated in Fig 3 C, for the same value of the stimulus *σ* = 0.3[1/*s*] and *τ* right *above* the critical point (*τ* = 1.7[*s*]), we can observe that accumulated evidence behind choices 1 and 2 fluctuates over time, which may account for perceptual switches. Furthermore, as shown in Fig 3 D, for large values of stimulus σ and the same value of the time delay, this effect disappears.

According to our extensive simulation of this model, presence of (1) values of delay above the critical point, for which the system crosses onto the unstable side; (2) a weak stimulus *σ*, are necessary for the stability switches to occur.

We then investigate the decision certainty in the function of the slope *β* and the stimulus magnitude *σ*. The results are presented in Fig 4. The model returns a monotonic, smooth and concave transfer function between both stimulus magnitude *σ* (Fig 4 B) and slope *β* (Fig 4 C), and the decision certainty. For certain combinations of parameters and for *t* high enough, this system can return firing rates *r*_1_(*t*), *r*_2_(*t*) below zero. In order to avoid such case we maximized certainty only on the time interval [0, 8].

**Figure 4.**
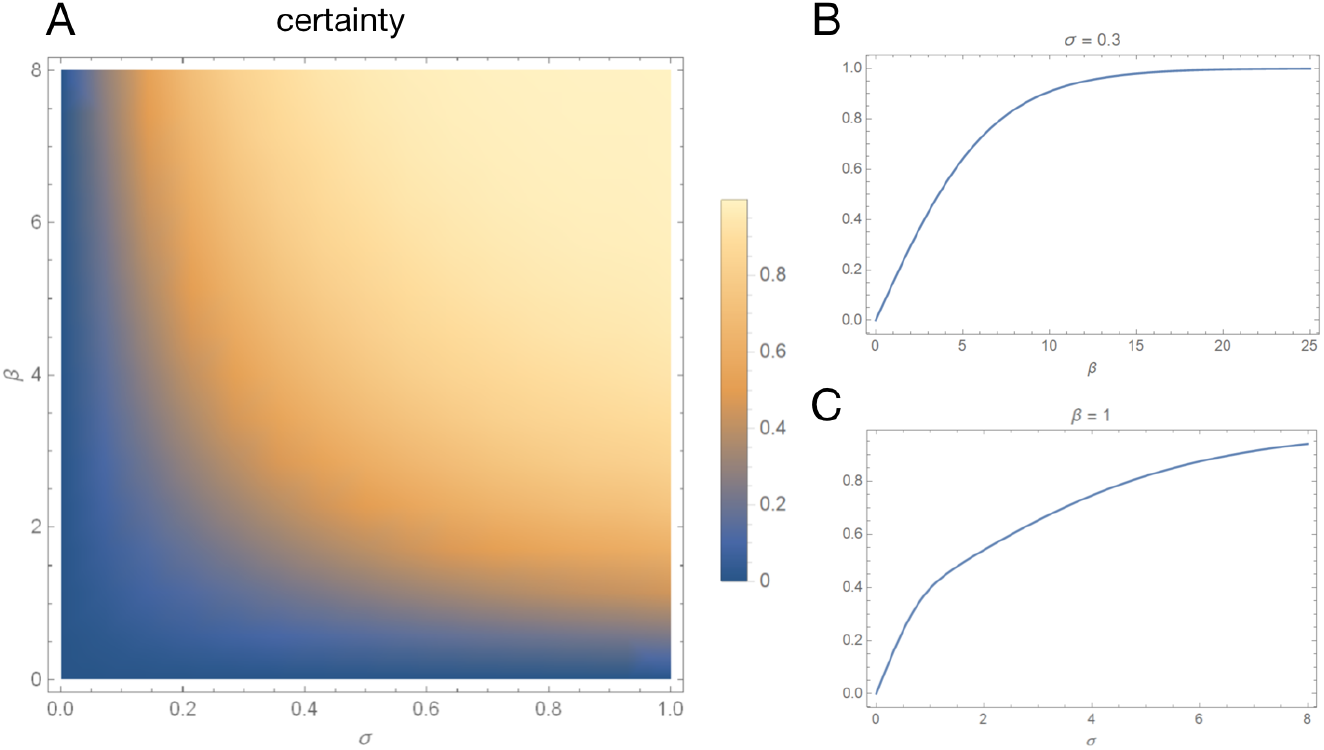
Features of the model. **A:** decision certainty in a function of slope *β* and stimulus magnitude *σ*. Certainty is a smooth and concave function of each one of the two parameters. B: certainty in a function of *σ*, for *β* = 1. **C:** certainty in a function of *β*, for *σ* = 0.3[1/*s*]. The model returns a monotonic, smooth and concave transfer function between both stimulus magnitude *σ* and slope *β*, and the decision certainty.

#### Influence of the stimulus duration on the decision making

One characteristic that our model exhibits, is that for a stimulus of certain magnitude (here, *σ* = 0.3[1/*s*]), perceptual ambivalence occurs for particular range of stimulus duration *t_max_* (here: for stimuli more brief than 1.0[*s*]), as presented in Fig 5.

**Figure 5.**
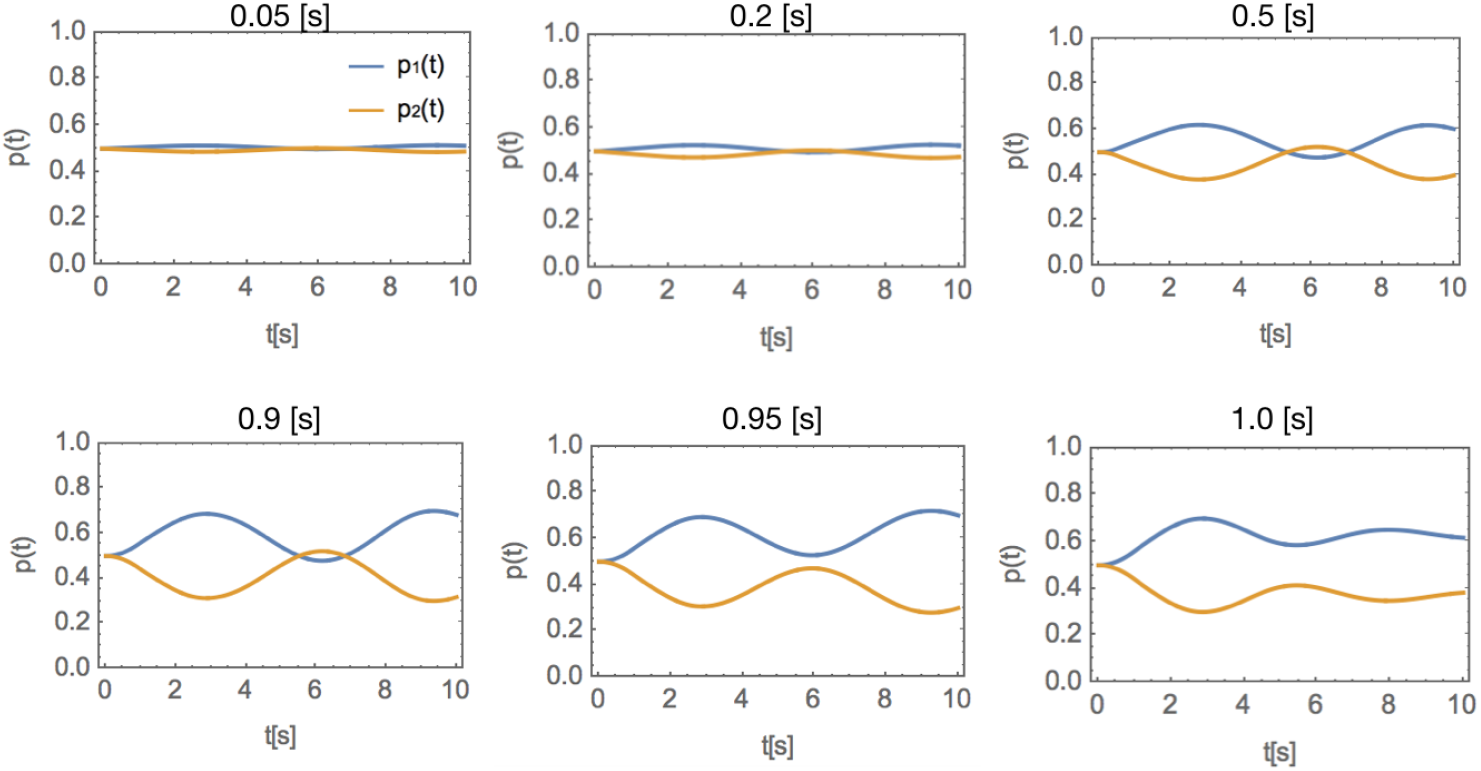
Dependency of the perceptual ambivalence effect on the stimulus duration *t_max_*. While the magnitude of the stimulus is fixed (*σ* = 0.3[1/*s*]), the duration of the stimulus is varying. For the stimuli longer than 1[*s*], the effect of ambivalence disappears.

#### Influence of the background noise on the decision making

In Fig 6, the dependency of the success rate on the magnitude of the noise is presented. The decision making is, in general, sensitive to noise, but the dependency of success rate in a function of noise magnitude is smooth as expected. Along with increasing *b*, the performance drops towards 0.5. Next to the results presented in Fig 6, we performed 20, 000 iterations of the system with *b* = 1.0, and the output success ratio was equal to 0.519 - which demonstrates that for large values of *b*, performance drops towards the chance level.

**Figure 6.**
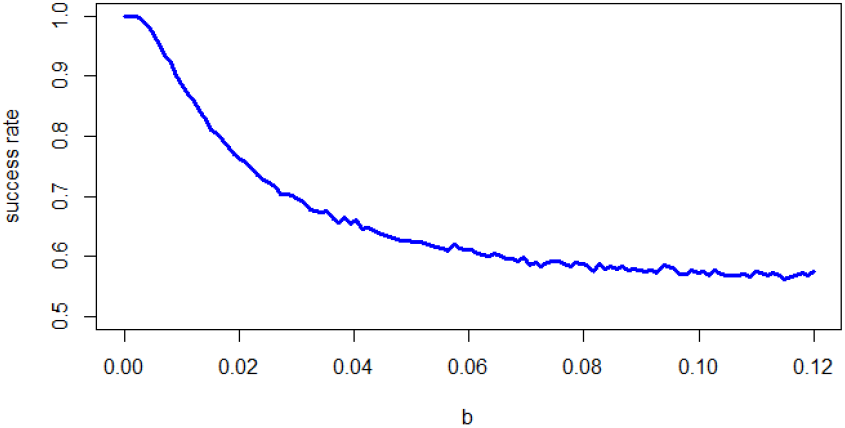
The dependency of the success rate on the magnitude of the noise generated from Wiener process.

### Validation in fMRI datasets

The variance in the three conditions within the SPL was equal to *var_w_* = 0.9148 and *var_s_* = 0.9720, respectively, and one-way ANOVA gives the statistic of *F* = 90.87 (*p* < *∊*). The two-sample, one-tailed post hoc F-test returns *var_w_* < *var_s_* at the significance level *p* = 0.0460. In the IPL, the associated variances were equal to *var_w_* = 0.9148, *var_s_* = 0.9720, respectively. The two-sample, one-tailed post hoc F-test returns *var_w_* < *var_s_* at *p* = 0.0050.

## Discussion

In this work, we bring together a few concepts popular in models of cortical networks. Local inhibitory networks are known as rhythm generators in the brain (both in the low [44] and high frequency range [45]). Furthermore, cortical networks are known to operate at the edge of instability [12]. These two aspects of the dynamics in cortical networks - sub-criticality and interneuron-induced rhythms - rarely come together in models of decision making. In this work, we propose a new population model which elicits the influence of dynamic state (close to the critical point) on the decision making process. We demonstrate that local, slow inhibition can induce Hopf bifurcation which activates distinct, context dependent modes of activity in the decision-making systems and can cause perceptual switches.

The two-population architecture resembles the model proposed by Ledoux & Brunel ([46], Fig 1) but in that work, one of the populations is excitatory while the other population is inhibitory. In our model on the contrary, the two populations are symmetric: both populations have local inhibitory and long-distance excitatory projections. Both network architectures can yield oscillatory behavior however, they should be applied to model different paradigms; architecture proposed by Ledoux & Brunel models behavior of one cortical population consisting of excitatory and inhibitory subpopulations (in response to time-dependent inputs) while our model models the competition between two distinct cortical populations.

As the model characterizes the role of local inhibition in decision-making, the neuronal delays play crucial role. We model these delays through Delayed Differential Equations (DDEs). DDEs are often used to model biological systems [20, 32, 47-54] because they exhibit a rich dynamical repertoire, and in particular, they fall into oscillatory regime on the edge of instability. DDEs were also used in the neural mass and neural field models of cortical activity [55-59]. Modeling the synapses with use of DDEs is not a common practice in computational relaxating variable instead of a delay. We chose for this type of modeling not in order to achieve a biologically plausible model, but to demonstrate a dynamical repertoire of networks with delayed synaptic transmission, similarly to work by Deco et al. [60]. As opposed to [60] however, we went beyond conditions for occurrence of the Hopf bifurcation, and we further investigated its role in the perceptual decision making.

Mathematically, both concepts for modeling synapses - delays in transmission and additional variable with relaxation - could be merged by the usage of distributed delay instead of discrete one. However, it is a common practice in modeling using DDEs that the first approximation is to use discrete-delay systems, as it is easier to analyze them both from mathematical and numerical point of view.

Various GABA receptors have various delays in the brain. I.e., the fast mode of inhibition is related to the GABA-A receptors [61, 62] which give synaptic delays lasting for several tens of milliseconds^3^. On the other hand, the slowest mode of inhibition is related to the metabotropic GABA-B receptors which have a time scale of a few hundred milliseconds [68]. Still little is known about the structure and functions of these receptors [69]. Finally, slow inhibition pathways [30], involving over 100 compounds, operate on timescales of hundreds of milliseconds to minutes, and is more complex than the synaptic transmission. In practice, the local inhibition in the nodes of the cortical network is most probably a combination of the multiple interacting processes at different time scales. In this work, we simplified the model to the very basic system with a single value of delay referring to slow inhibition processes, as this work is a proof of concept.

In general, the inter-population excitatory connectivity can also be delayed (especially the glutamatergic NMDA receptors which give the highest delays). However, mathematically, these delays in excitation do not generate oscillations on their own and therefore do not change the dynamic properties of the model, therefore we skipped these delays from the model for the sake of simplicity.

We also used a simple form of a stimulus. We chose as a step impulse of the duration *τ*. DDEs have an infinite dimensional space of initial conditions as the initial condition on the interval [–*τ*, 0] can be defined as any continuous function. This opens a range of possibilities for the future research on the dynamical properties of the decision making systems with delays, for instance in a function of the stimulus properties. With respect to the stimulus magnitude, we need to remember that the model is a firing rate model, therefore the weak input of *σ* = 0.3[1/*s*] corresponds to a higher firing rate in the upstream populations projecting to the system. Typically, connectivities in cortical networks are modeled as having the density of 10%, which corresponds to the firing on the speed of 3.0[1/*s*], which equals 750 % of the background firing in the population model (*I*(*t*) = 0.4[1/*s*]). For the strong stimulus, *σ* = 1.0[1/*s*] corresponds to the firing rate of 10.0[1/*s*] in the upstream populations, which is 2500 % of the background rate, consistent with experimental results [70]. On the other hand, the unconstrained firing predicted by the model in case of strong stimuli is not in concordance with experimental evidence: transfer functions in neurons tend to be sigmoidal [71], therefore the activity should saturate for strong stimuli.

Furthermore, as the self inhibition rather than synaptic plasticity is the crucial feature in the model, we chose for the simple form of plasticity: a sigmoidal function of the multiplication between *r*_1_(*t*) and *r*_2_(*t*) - which is a simple implementation of the Hebbian rule. We included synaptic plasticity because it is the integral part of learning and therefore also of the decision making [72]. However synaptic plasticity is not central mechanism in this model as it does not affect the dynamical repertoire of the system.

The resulting model has interesting properties. The certainty of the decision making is a smooth, monotonic function of both the stimulus magnitude *σ* and the slope *β* (Fig 4 A), and saturates at 1. The *β* parameter denotes the slope which quantifies the influence of the difference between *r*_1_(*t*) and *r*_2_(*t*) on the difference in probabilities *p*_1_(*t*) and *p*_2_(*t*). The monotonically increasing function of certainty in the function of *β* is one of the assumptions in the model (Eq. 4, Fig 4 B). On the other hand, the monotonic, increasing function of certainty with respect to *σ* on the other hand is not an assumption, but an emergent *feature* of the model (Fig 4 C), and is consistent with expectations: the ambivalence in decision making disappears once the stimulus becomes strong enough.

Furthermore, we also found that, for a stimulus of certain magnitude, perceptual ambivalence occurs for particular range of stimulus duration (Fig 5). Although we did not find any literature reporting experimental findings on the influence of the stimulus duration on perceptual bistability, we believe that this effect is concordant with common sense, as presenting a stimulus for a prolonged period of time should allow for collecting more evidence on behalf of one option over the other.

There is a variety of neuroimaging techniques which can be used to validate this model. We chose to validate the model with fMRI datasets, which were also previously employed to studies on perceptual decision making [73]. The temporal resolution of fMRI recordings is very low. Therefore, we attempt to overcome this issue by operationalizing the problem as the variance of the joint signal. Note that the expected value of the variance is indifferent from the TR of the fMRI sampling, and the accuracy of its estimation only depends on the length of the signal. Furthermore, since the direct location of the competing populations in the cortex cannot be established, we incorporated the two parietal areas previously reported to be involved in solving the perceptual ambiguity into the study, SPL and IFL, in their entirety. As neuronal oscillations are notoriously hard to capture in the fMRI experiments, we based the hypothesis on the *variance of the joint signal* coming from the competing populations. Given such a simplification, the nearby cortical populations oscillating in anti-phase, the joint signal collected from these populations should drop in variance^4^. Given these assumptions, we found that the variance of the BOLD signal indeed drops in SPL and IPL on the group level after presentation of ambiguous stimuli as compared to the variance in absence of the stimuli, or in presence of strong, unambiguous stimuli. This is, of course, only one clue behind the bifurcation model of decision making introduced in this work, and more in depth neurophysiological validation is necessary.

An alternative validation method could be electroencephalography (EEG): the oscillations in anti-phase will yield higher power than the oscillations in the in-phase and, as such, they should be detectable from the EEG readout. As we do not have any extra knowledge upon the detailed mechanisms underlying facial perception, and as our literature-informed regions of interest are relatively small, we believe that in this particular case, fMRI is a better validation technique than EEG. In the future, it might be possible to use also EEG activity transformed into the cortical space through the inverse problem, and use power spectrum for validation of the model.

We chose the task on the perception of gender-ambiguous faces for three major reasons. Firstly, there are exactly two possible choices in this task as the cortical networks need to disambiguate the gender of the presented face. Secondly, the face recognition is a natural, evolutionary and involuntary mechanism [74], therefore it naturally engages the brain and is associated with a natural motivation to disambiguate the stimulus, regardless of the level of external motivation, or fatigue in a participant. Thirdly, in this experiment faces are presented for a relatively short period of time (0.5[s]), followed by free viewing of the screen. This setup corresponds to our model, in which we are simulating the phase transition in the dynamics of the system in a reaction to a stimulus presented to one of the populations for 0.5[s].

In the future research, the model should also be validated with respect to other experimental paradigms related to perceptual bistability: visual (such as the Necker cube [1]) or auditory (such as mixtures of high and low tones, which can be interpreted by the brain in multiple ways [75]).

In this work, we present a proof of concept that the slow inhibition can yield a rich dynamics in the decision-making systems. This model has a few implications. Firstly, perceptual switches present while watching ambiguous stimuli (such as the Necker cube) are often envisaged as a result of the noise driven switches between attractors in the decision-making systems [76]. In our model however, the perceptual switches can arise even in absence of the noise, as a natural consequence of stimulation of the systems with delayed self-inhibition.

## Author Contributions

KP, MP, UF and NB designed the model. PR, NB, KP and MP carried out the initial simulations. PR write Mathematica implementation of the model, and implemented R codes. LT designed the pipeline and conducted the fMRI experiment. UF and NB critically reviewed the results. NB, UF and LT wrapped the body of the manuscript.

## Acknowledgments

We would like to thank **Maciej Borodzik, Piotr Kozakowski, Bard Ermentrout, Andrzej Kozłowski, Piotr Suffczyński** and **Franciszek Rakowski** for the discussion and advice. We would also like to cordially thank to **Jan Poleszczuk** for advice and fruitful brainstorming sessions.

## Supporting information

### S1 Stability analysis of the model

Consider the system of equations 
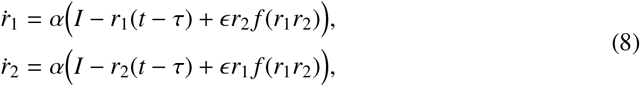

where 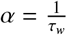 is a time scale, *I* is a constant input and 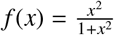. We are mainly interested in the model dynamics for reference parameter values

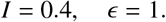

On the other hand, as the qualitative dynamics of Eqs. (8) does not depend on the magnitude of *α*, in the following we assume *α* = 1.

Looking for steady states we obtain

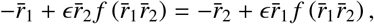

where (*r*̄_1_, *r*̄_2_) denotes the steady state. This relation implies

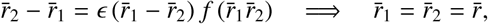

as *f* (*x*) ≠ −1/*∊* due to non-negativity of this function. Therefore, the coordinate *r*̄ satisfies

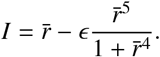

Let us consider an auxiliary function defined as 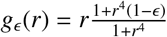, and for the reference *∊* = 1 this function reduces to 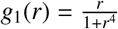. It is easy to see that *g*_1_ is unimodal, *g*_1_(0) = 0, 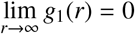, and has its maximal value 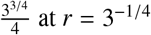. This means that there are at most two steady states, depending on the value of *I* > 0. On the other hand, if *∊* ≠ 1, then the derivative 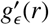 could have two positive zeros, and therefore there are at most three steady states. For the reference *I* = 0.4, let us consider another auxiliary function 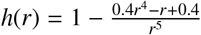. There is 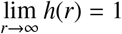 and 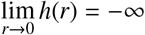. Moreover, *h*(*r*) > 0 for *r* > 0.4 and we can check that *h’*(*r*) has two zeros in the interval (0, 2), around 0.5065858562 (for which there is maximum of *h* around 3.405131243) and 1.952080163 (for which there is minimum of *h* around 0.84984565). Hence, there are at most three steady states, depending on the magnitude of *∊*.

#### Corollary 0.1.

*If ∊* = 1*, then*

- *for 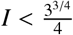 there are two steady states of* *Eqs.*(8);
- *for 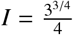 there is exactly one steady state of* *Eqs.*(8);
- *for 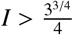 there is no steady state of* *Eqs.*(8).

*If I* = 0.4*, then*

- *for ∊* < 0.84984565 *and ∊* ≈ 3.405131243 *there is exactly one steady state of* *Eqs.*(8);
- *for ∊* ≈ 0.84984565 *and* 1 ≤ *∊* < 3.40513124 *there are two steady states of* *Eqs.*(8);
- *for* 0.84984565 < *∊* < 1 *there are three steady states of* *Eqs.*(8);
- *for ∊* > 3.405131243 *there is no steady state of* *Eqs.*(8).

#### Remark 0.2.

*For the reference parameter values ∊* = 1 *and I* = 0.4 *there are two steady states of* *Eqs.*(8)*, around* 0.4114655 *and* 1.1827404.

Looking for local stability of these states for *τ* = 0 we calculate Jacobi matrix for Eqs. (8) obtaining

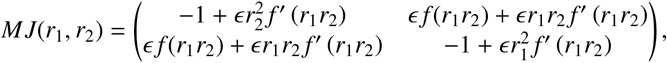

and for a steady state (*r*̄, *r*̄) it reads

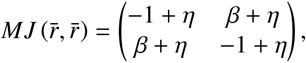

where *η* = *∊r*̄^2^ *f’* (*r*̄^2^ > and *β* = *∊ f* (*r*^2^ > 0.

We have tr *MJ* (*r*̄, *r*̄) = −2 + 2*η* and d_(_et *MJ* (*r*̄, *r*̄) = 1 − 2*η* − 2*ηβ* − *β*^2^. Hence, the necessary condition of stability is *η* < 1, that is 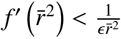. Moreover, calculating the determinant of the characteristic equation we easily see that it is always positive, such that any steady state is either a saddle or a node.

For *r*̄ ≈ 0.4114655 we have det *MJ* (*r*̄, *r*̄) ≈ 0.7862262548 and tr *MJ* (*r*̄, *r*̄) ≈ −1.891645542, which means that this point is a stable node. For *r*̄ ≈ 1.1827404 we have det *MJ* (*r*̄, *r*̄) ≈ −0.5901784196, which means that this state is a saddle.

#### Corollary 0.3.

*For the reference parameter values I* = 0.4 *and ∊* = 1 *the steady state with smaller coordinates is a stable node, while the steady state with greater coordinates is a saddle.*

Moreover, analyzing the phase space portrait we are able to check that all solutions below the stable manifold of the saddle are attracted by the stable node, while above this manifold all solutions go to infinity.

Now we turn to the case *τ* > 0. We know that if a steady state is a saddle for *τ* = 0 then it remains unstable for all *τ* > 0. Hence, we want to check if stability switches are possible for the state (*r∊*, *r∊*), *r∊* ≈ 0.4114655. Calculating the characteristic matrix one gets

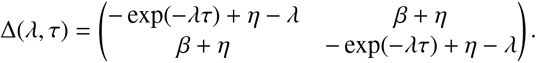

To get stability switches one needs to find eigenvalues λ = ±*iω*, *ω* > 0, in the imaginary axis and check if they cross this axis.

Let us take *λ* = *iω*. Then

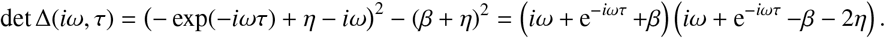

It is obvious that det ∆(*iω*, *τ*) = 0 iff *W*_1_(*iω*, *τ*) = *iω* + e^−*iωτ*^ +*β* = 0 or *W*_2_(*iω*, *τ*) = *iω* + e^−*iωτ*^ –*β* –2*η* = 0. As *W*_1_ and *W*_2_ are very well know transcendental equations, we can conclude that

1. if *β* + 2*η* < 1, then there are two sequences of critical delays 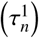 for *W*_1_ and 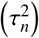 for *W*_2_ at which stability switches are possible;
2. if *β* < 1 and *β* + 2*η* > 1, then there is only one sequence of critical delays corresponding to *W*_1_;
3. if *β* > 1, then stability switches are not possible.

It occurs that for the reference parameter values *I* = 0.4 and *∊* = 1 the first case occurs. Moreover, we know that for the first critical delay the stable steady state loses stability and cannot gain it again. This means that we can suspect that solutions of Eqs. (8) oscillates permanently for sufficiently large delays.

### S2 The Balloon-Windkessel model of the hemodynamic response function

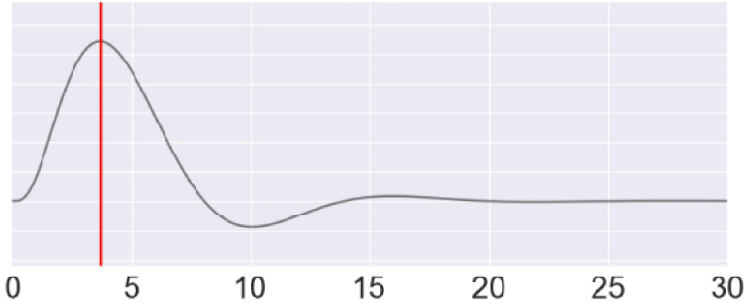
The canonical HRF response (κ = 0.65[1/*s*], *γ* = 0.41[1/*s*], *λ* = 0.98[*s*], *α* = 0.32, *ρ* = 0.34)

The classic Balloon-Windkessel model for the hemodynamic response [34, 36] reads as follows

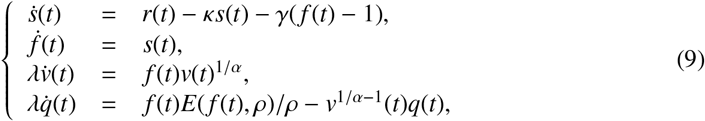

where *r*(*t*) – the underlying, fast neuronal dynamics, *s*(*t*) – vasodilatory signal, *f* (*t*) – inflow, *v*(*t*) – blood volume, *q*(*t*) – deoxyhemoglobin content, *E*(*f*, *ρ*) = 1 – (1 – *ρ*)^1/ *f*^. This model contains five node-specific constants: *κ* – rate of signal decay, *γ* – rate of flow-dependent elimination, *λ* – hemodynamic transit time, *α* – Grubb’s exponent, *ρ* – resting oxygen extraction fraction.

Then, the following expression describes the BOLD response

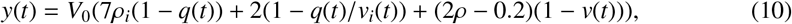

where *V*_0_ = 0.02 is the resting blood volume fraction. The hemodynamic parameters were set at the mean of the distributions given in [34]: *κ* = 0.65[1/*s*], *γ* = 0.41[1/*s*], *λ* = 0.98[*s*], *α* = 0.32, *ρ* = 0.34.

This generative model characterizes BOLD response *y*(*t*) in a function of fast neuronal dynamics *r*(*t*) as a system of ordinary differential equations. Effectively, *y*(*t*) can also be obtained from *r*(*t*) as a linear convolution with a kernel which describes BOLD response to a pulse stimulus *r*(*t*), also known as a hemodynamic response function (Fig 7).

## Footnotes

1 Furthermore, testing the role of the frontal lobe activity in paradigms involving bistable stimuli (binocular rivalry or bistable perception) is difficult as it can be confounded with modulations of attention [26] or reflect its role for conscious visual perception [26, 27]

2 This relaxation term is often used in modeling neuronal activity, from modeling Poissonian spike trains of single neurons [33] to modeling interacting populations in Dynamic Causal Modeling [34]

3 excluding the afterdepolarizations lasting for several tens of milliseconds as well [63-67]

4 We assumed that, even if a part of the anatomically defined ROIs is not involved in this particular paradigm, this will not affect the results, because it will contribute to the variance across all three conditions in the same fashion

